# Clustering of plasmid genomes for genomic epidemiology by using rearrangement distances, with pling

**DOI:** 10.1101/2025.09.02.673752

**Authors:** Daria Frolova, Zamin Iqbal

## Abstract

Integration of plasmids into genomic epidemiology is challenging, because there are no clearly defined evolving-units (equivalent to species), and because plasmids appear to evolve as much by structural change (rearrangements, insertions and deletions) as by mutation (1). Further, plasmids transfer horizontally between bacterial hosts (2), and thus a model beyond just a phylogeny is needed to integrate their genetic information with that of their hosts.

Pling (3) is a tool designed to measure a genetic distance between plasmids that is related to how they empirically appear to evolve, by measuring the distance between two plasmids as the minimum number of structural changes needed to change one plasmid into the other (ignoring SNP differences). Having done this, it constructs a relatedness network of the plasmids under study, and then clusters them into groups that are credibly recently related.

We give here a protocol for running pling, and how we integrate its information with plasmid typing and SNP information. Together, these provide a system for deciding which plasmids are worth treating as “the same plasmid” for the purposes of epidemiology, quantifying their relatedness in terms of rearrangements and SNPs, and then seeing how they are distributed across the host phylogeny.

## Introduction

The use of whole genome sequence data to study the spread and evolution of bacterial pathogens is now routine, and there are robust computational and modelling methodologies for inferring relatedness between genomes (4–9). These benefit from the fact that there is a good model of bacterial evolution - namely, the phylogeny - which captures the clonal frame and speciation, although it is blind to horizontal gene transfer and recombination. Second, that it is often practically useful to be able to cluster genomes as “the same” when a specific timescale is being considered. Commonly this comes with an associated set of nomenclatures for “types” (10–14).

Together, these provide the key tools which are applied to the study of bacterial outbreaks in public health. When considering plasmids, the first problem is that we cannot easily group them into “species” for independent analysis, and the second is that there is no underlying evolutionary model to which we can turn. Plasmids evolve through multiple independent processes – mutation at replication time (15), apparent mutation through “classical” recombination (16, 17), gene gain and loss (18, 19), inversions (20), and mobile-element mediated rearrangements (21). Here, it appears we cannot focus first on SNPs and ignore the others as “second-order issues”.

Historically there have been various approaches taken for plasmid classification (see (22) and references therein) initially involving typing based on gene alleles, and more recently using whole plasmid genome comparisons. These latest approaches involved kmer-based estimation of ANI using mash (or similar) tools (23–25), creation of a relatedness network where edge weight represented mash distance, and then some kind of clustering on the network (26–28). This approach allows a data-driven method of discovering “natural” groupings of plasmids. However there are challenges. First, by using mash (or equivalent tools) to estimate ANI, one effectively samples a fixed number of kmers to compare genomes - this works well for bacteria as all the genomes are approximately the same size, but could “penalise” events in which a significant proportion of a plasmid is gained/lost in one step. Second, it would be good to have a genetic distance where the distance scales with number of events, not k-mers. Third, transposons/IS elements which are shared between unrelated plasmids can create a false signal of relatedness if one simply looks for shared sequence. These problems were addressed in detail in (3) which introduced two innovations. First, it used two genetic distance measures, one (called “containment distance”) which measures how much of the smaller plasmid is contained in the larger, and the other (DCJ-Indel (29)) which counts the number of rearrangement events (gain/loss of sequence, inversion, transposition) between two plasmids. Second, it made a relatedness network based on the containment distance and DCJ-Indel distance, and first identified likely transposons from the network topology (as they connect unrelated plasmids), before finally clustering. This allowed pling to identify plasmids that are related through structural events to be clustered together, while avoiding transposon-drive noise. The paper showed how this resulted in larger core genomes within clusters.

In this paper we detail a protocol for running and interpreting pling.

## Materials

Pling is a conda installable command line tool and runs on Linux. It can be run on a laptop or on a compute cluster. Using sourmash (30) prefiltering, 300 plasmids can be clustered on a laptop in about 20 minutes. If running on >1000, we would recommend using a cluster.

## Method

### 1. Pling approach

Pling calculates containment and DCJ-Indel distances and uses these to construct a network (with plasmids as nodes) and then clusters plasmids on the basis of this. The DCJ-Indel distance is based on the “double cut and join” operation – you cut at two positions in the genome, and then join the four ends in a new way (Figure 1). The distance is then the minimum number of DCJ and insertion or deletion operations necessary to transform one genome into another (29, 31, 32). It is defined on ordered integer sequences, in which integers represent blocks of sequence. In pling, these integer sequences are generated by aligning pairs of plasmids to each other, and giving matches the same integer marker (default minimal sequence similarity 80%), while unmatched sequences (of least 200 bases with default settings) are given unique markers. This alignment is also used to calculate the containment distance between two plasmids, i.e. what proportion of the smaller plasmid is not contained in the larger.

**Figure 1:**
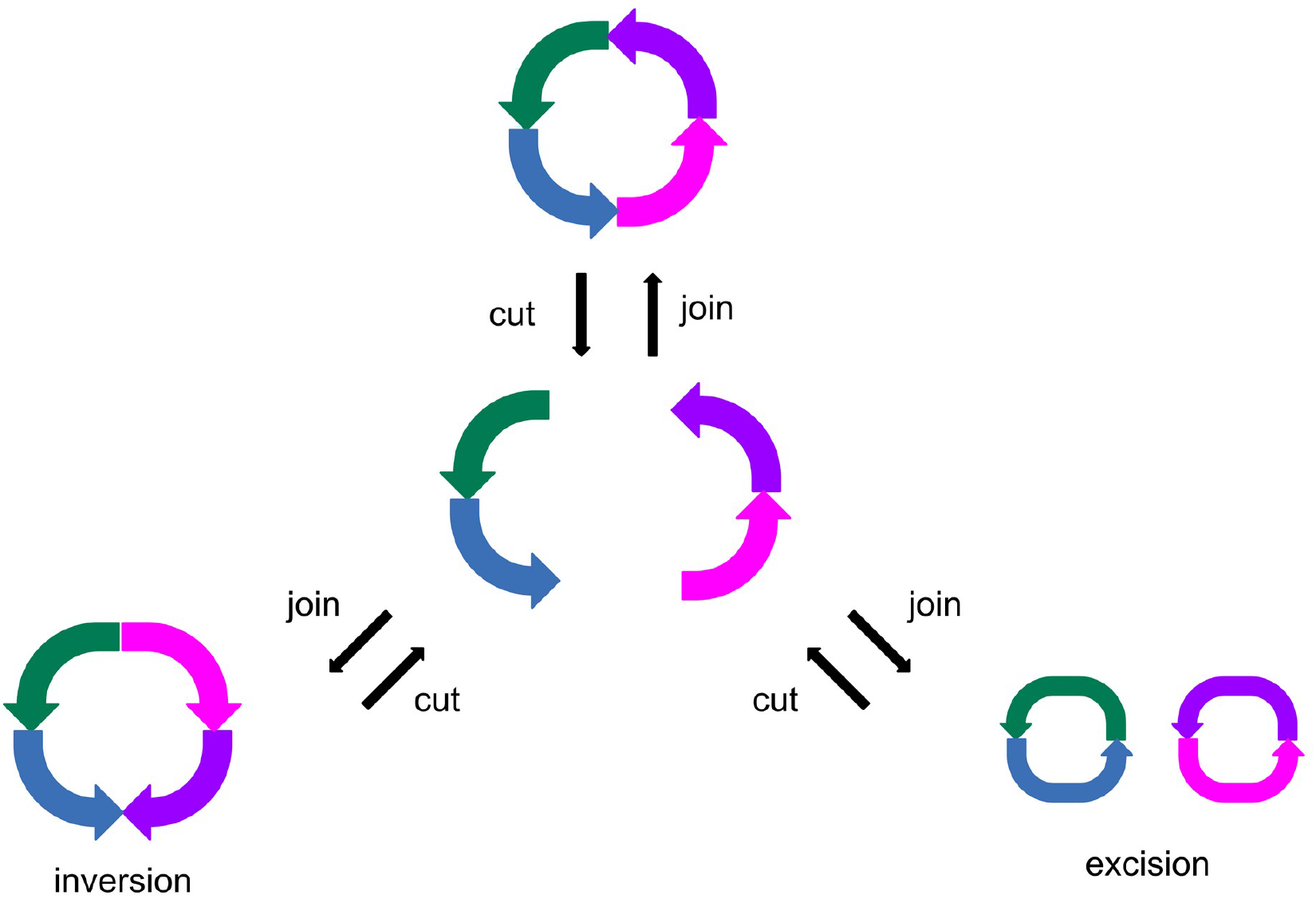
The DCJ operation on a circular plasmid. Each coloured arrow represents a marker. On a circular plasmid, a double cut and join operation will create either an inversion, or an excision. Each operation can be reversed by cutting in the same positions, and then rejoining appropriately.

The containment distance is used as a first approximation of relatedness between plasmids. This is done by constructing a containment network: nodes represent plasmids, and plasmid pairs with containment beneath a given threshold (default 0.5) are joined by an edge. Connected components on this network define plasmid *communities* – these are not meant to be used for typing, but rather to place plasmid types within a broader context.

Plasmids connected by an edge in this network will then have the DCJ-Indel distances calculated between them, and then a subnetwork is induced on the containment network by removing any edges between plasmids with DCJ-Indel distance above the DCJ-Indel threshold (default 4). We refer to this subnetwork as the DCJ-Indel network.

At this point, pling looks for hub plasmids, which are plasmids that are densely connected, but their neighbours are sparsely connected to each other, indicating that these plasmids are not truly related to the hub or each other. The motivating example for hubs is a transposon which is shared by unrelated plasmids. With default parameters, nodes are labelled as hubs if they have at least 10 neighbours, and those neighbours have an edge density less than 0.2. The edge density is the ratio of the number of actual edges to the number of all possible edges. To prevent these hub plasmids from impacting the clustering, they are removed from the network. This is then finally what we refer to as the pling network, and this is the network used to cluster on. There is a toy example of this network in Figure 2. The clusters found on the pling network are called *subcommunities*, and these determine what type a plasmid is. Hence, every plasmid (that is not a hub) will be part of a containment community, and a DCJ-Indel subcommunity.

**Figure 2:**
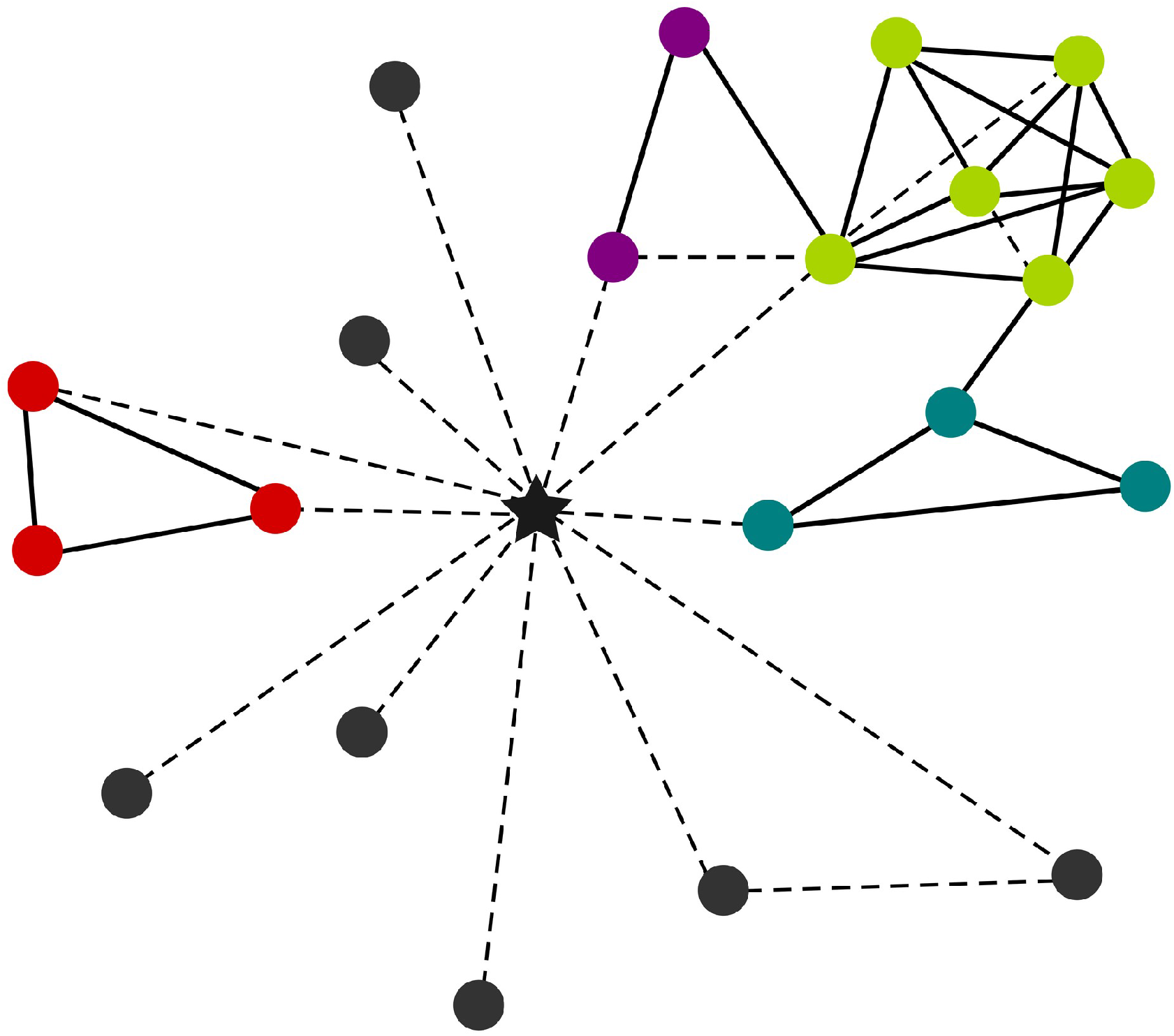
Toy pling network. Each node represents a plasmid, and together with solid edges show a toy pling network. The dashed lines represent edges that were removed either due to the plasmid pair being over the DCJ-Indel threshold, or due to being labelled hub (here denoted by a black star). This network overall shows a *community* of plasmids, while the colour of the nodes denotes the *subcommunity*. Red, purple, green and blue nodes are each distinct subcommunities, while the grey nodes are singletons.

### 2. Installation of pling

Pling is installable via conda with the following commands:

> *conda config --add channels bioconda*
>
> *conda config --add channels conda-forge*
>
> *conda install pling*

Alternatively, pling can be cloned directly from GitHub, via

> *git clone https://github.com/iqbal-lab-org/pling.git*

The dependencies can be found in the *env*.*yaml* file, which can be used directly to create a conda environment. Finally, pling will need to be installed within its directory with

> *python3 -m pip install*.

Installation through Docker is also possible.

Pling has an optional dependency, Gurobi (33). This is used when calculating DCJ-Indel distances, and is faster than the default tool used, which is GLPK (34). Gurobi is a commercial software that requires a licence, which is free for academic use and can be obtained from their website. Version 10.0.1 of Gurobi can then be installed through conda, using the channel https://conda.anaconda.org/gurobi.

### 3. Other tools

Supplementary information gained from other tools can be very useful when working with pling and its output. We suggest MOB-suite (28) for general metadata about plasmids, such as Inc types and mobility. For pangenome analyses, we use ggcaller (35) and pangraph (36), while for creating SNP trees we recommend parsnp (37).

All these tools are available on GitHub, either as downloadable binaries or with installation through conda or Docker.

### 4. Data preparation

Pling requires individual fasta files for each plasmid to run, and we recommend that each file is named after the plasmid ID. They do not need to be all in the same directory. The plasmid genomes should be complete, but do not necessarily have to be circular, although pling will assume they are circular by default. The input to pling is a text file consisting of a list of file paths to each fasta.

### 5. Run pling

To run pling with defaults, just run:

> *pling input*.*txt output_dir align*

where input.txt is the input text file with a list of fasta filenames, and output_dir is the name of the output directory.

If running on a large dataset, you will likely want to run with multiple cores/threads, increased batch sizes, and using the sourmash prefiltering. ‘Batch size’ refers to batches of pairs of plasmids, because pling needs to compare all plasmids against all; the pairwise comparisons are split into batches, to enable parallelisation. The sourmash prefiltering removes plasmid pairs with containment distance greater than 0.85 early in the workflow, which reduces runtime, however there are sometimes issues using it for very small plasmids (∼5kb or less). You can do this by, for example, running

> *pling input*.*txt output_dir align --cores 8 --batch_size 1000 --sourmash*

You can change the prefiltering threshold using the *--sourmash_threshold* argument, for example if you want to filter out more pairs, you could choose a threshold of 0.65. When changing the sourmash threshold, it is important to remember that sourmash is prone to overestimate the containment distance, so to avoid wrongly filtering out pairs, we recommend setting the sourmash threshold 0.25 or 0.35 higher than your overall containment threshold.

If you wish to include some metadata in the visualisations, you can feed a tsv of metadata to pling via *--plasmid_metadata*. This metadata will be added beside the plasmid ID on the visualisation network. Note that this is only for adding one piece of additional metadata. Good choices are replicon types, genome lengths, or predicted mobility – these can be easily generated by running the *mobtyper* command from MOB-suite.

To run pling on a cluster, such that each batch of computations is submitted as an individual job, you need a cluster profile – luckily, these can be easily created from the recipes found in (38). You can then set it to run on a cluster, for example, like this

> *pling input*.*txt output_dir align --profile cluster --batch_size 1000 --sourmash*

where *cluster* is the name you gave to your cluster profile.

### 6. Pling output and viewing network visualisations

The output consists of 4 folders (named *batches, containment, dcj_thresh_4_graph, unimogs*) and one tsv file (named *all_plasmids_distances*.*tsv*). Generally, you will only need the tsv file *all_plasmids_distances*.*tsv* and the folder *dcj_thresh_4_graph*, so we focus on these here. The file contains the DCJ-Indel distances for all plasmid pairs that pass the containment distance threshold, while the folder contains the final pling typing and the pling network, including visualisations. In *dcj_thresh_4_graph*, there are two subfolders named objects and visualisations. In objects are the files for the pling typing (*typing*.*tsv*) and a list of hub plasmids (*hub_plasmids*.*csv*). In visualisations there are also two subfolders, named *communities* and *subcommunities*. Each of these subfolders has the same structure: a file named *index*.*html*, and two folders named *graphs* and *libs*. These three objects need to be kept together in one folder for the visualisations to work.

Briefly, the remaining folders (*batches, containment* and *unimogs*) contain outputs from intermediary steps of the pling workflow; *containment* contains a file of containment distances and containment network visualisations, *unimogs* is the output of the integerisation step, while *batches* is used internally and is primarily retained for debugging purposes. If you run with sourmash prefiltering, there will also be a fifth folder *sourmash*, which contains the results of running sourmash.

To view the community visualisations, go to *dcj_thresh_4_graph/visualisations/communities*, and open *index*.*html* with your browser of choice (although in our experience, on Windows you need to use Chrome). This is an index of all the plasmid communities, and it should look similar to the example in Figure 1. You can view the network for any community by clicking on the links in the index. In the network visualised, each node is a plasmid and is coloured by which subcommunity the plasmid belongs to (note that colours can repeat, but the visualisation avoids placing repeating colours next to each other). Black stars denote hub plasmids. There is an edge between two plasmids if they pass the containment threshold only – this is therefore a visualisation of the containment network, and not the DCJ-Indel network, and includes edges to hubs. An edge is labelled with the containment and DCJ-Indel distances (containment / DCJ-Indel) for the pair of plasmids it connects. You can click on nodes and move them around to change the layout of the network; the default layout groups nodes together based on how densely connected they are, without accounting for distances. For subcommunities the visualisation works the same, but each entry in the index is for a subcommunity, and it shows edges for pairs of plasmids that pass the containment and DCJ-Indel thresholds, i.e. it visualises the DCJ-Indel network. The colouring of nodes is consistent across both community- and subcommunity-level visualisations, allowing for easy comparison. For example, in Figure 3 the bright green subcommunity can be seen in the left hand side corner of the community visualisation.

**Figure 3:**
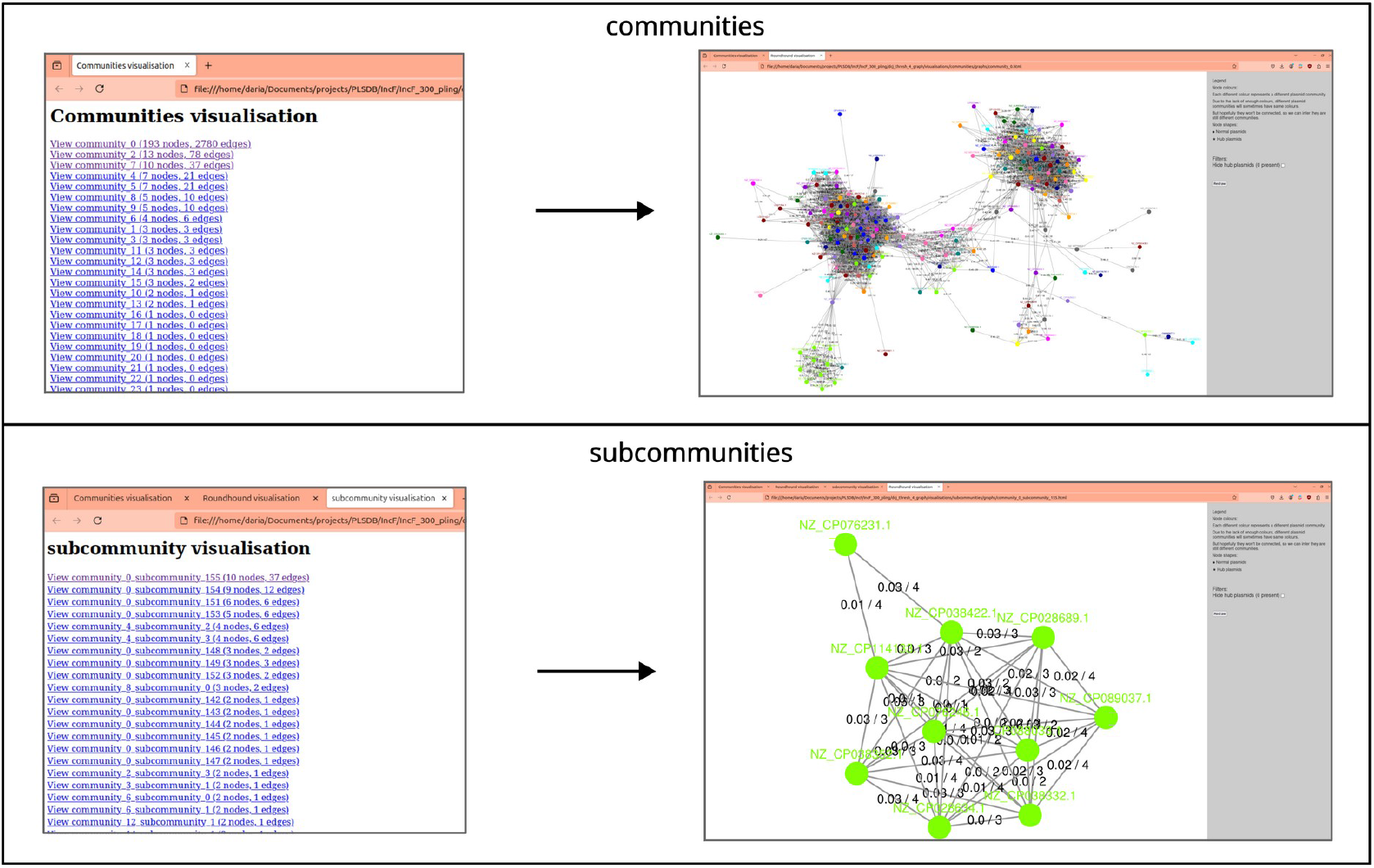
Examples of how the visualisations look for pling communities and subcommunities. The upper panel shows what to expect the index and network visualisation to look like for communities – when opening the index, it opens to a page of links to containment community networks, ordered by size. The lower panel shows the same for subcommunities, but these visualise DCJ-Indel networks for each subcommunity instead. This network was created from IncF plasmids from PLSDB (45). A list of these plasmids can be found in (40).

By default, pling only outputs these html visualisations, but running pling with *--output_type both* will produce Cytoscape-formatted json files for each community and subcommunity, which can be loaded directly into Cytoscape. They can also be used with some R and python packages for graphs.

### 7. Assessing quality of the clustering

There are several characteristics by which you can evaluate whether a pling clustering is satisfactory. The most effective is to calculate the core genome and check its median relative size for each subcommunity – this is the metric we used in (3). The relative core genome size is found by summing the length of the core genes present in a plasmid, and dividing by the total length of the plasmid. Note that for plasmids, often a less strict threshold is chosen to define the core; for example, presence in 80% of genomes has previously been used (39).

We suggest using ggcaller for calculating core genes in a subcommunity, since it is very fast on plasmids, and computes both gene annotations and the pangenome simultaneously. The documentation for it is available on GitHub, and we run it with default settings. For a script to parse the output for core genes and calculate the median, you can base it on the script in *relative_core_genome* in (40).

Alternatively, pangraph can be used to calculate core “blocks”, rather than genes. Pangraph is a pangenome tool that segments a group of genomes into shared and indel synteny blocks, and has a convenient Python library for parsing its output. However, its definition of a core block excludes duplicated blocks, so in subcommunities with many duplication events, it can underestimate the core genome.

Another metric is the distribution of plasmid lengths within a subcommunity, which can be evaluated using a histogram of bin counts. If the distribution is tight and looks approximately normal, then this is a good sign. Some subcommunities may have bimodal length distributions – this is not necessarily an indicator of the clustering being poor, as this can happen when there have been fusion events in the subcommunity. It does warrant verifying that the cluster is sensible though, and this can be done by viewing the network, which we discuss how to interpret further below.

Finally, checking the replicon types of subcommunities is also a useful measure, and can easily be done by running MOB-suite. Especially in the case of fusions, there may be mixed replicon types in a subcommunity; however, you would typically expect the clustering to be mostly compatible with replicon typing. Additionally, untyped plasmids will often not only form a separate subcommunity, but even a separate community.

If there appear to be any issues with the clustering, then thresholds may need to be adjusted, which we discuss in the next section.

### 8. Changing thresholds

The containment distance threshold and DCJ-Indel distance thresholds can be modified through the arguments *--containment_distance* and *--dcj*. The default thresholds are 0.5 and 4 respectively, but for example if you wanted to cluster plasmids with a lot of sequence similarity but more structural variation, you could use the command:

> *pling input*.*txt output_dir align --containment_distance 0*.*1 --dcj 7*

If you want to try multiple DCJ-Indel thresholds, you can rerun pling with the same output directory, and it will reuse previous results and produce a new folder named (if, say, your new threshold is 7) *dcj_thresh_7_graph* in the same output directory. However, changing the containment threshold triggers rerunning the whole workflow, so make sure to use a different output directory.

The difficulty lies in deciding when changing thresholds is useful. While generally the defaults work well, they may not always be suited to your specific use case. For example, you may find that the clustering is too fine and is separating plasmids that you want to be grouped together. In Figure 4 is the visualisation of a community network constructed with default thresholds, which has lilac, turquoise and green subcommunities only separated by DCJ-Indel distances 5-8, and containment distances less than 0.1; but distances to the remaining plasmids are at least 10 DCJ-Indel, and they have larger containment distances. Here it would make sense to change the DCJ-Indel threshold to 8, thus joining the three subcommunities into one. We would not typically recommend setting the threshold any higher than 10 though, as our experience is that at DCJ-Indel distances of 10 result in alignments between plasmids that are very complex, and unlikely to reflect a recent relationship.

**Figure 4:**
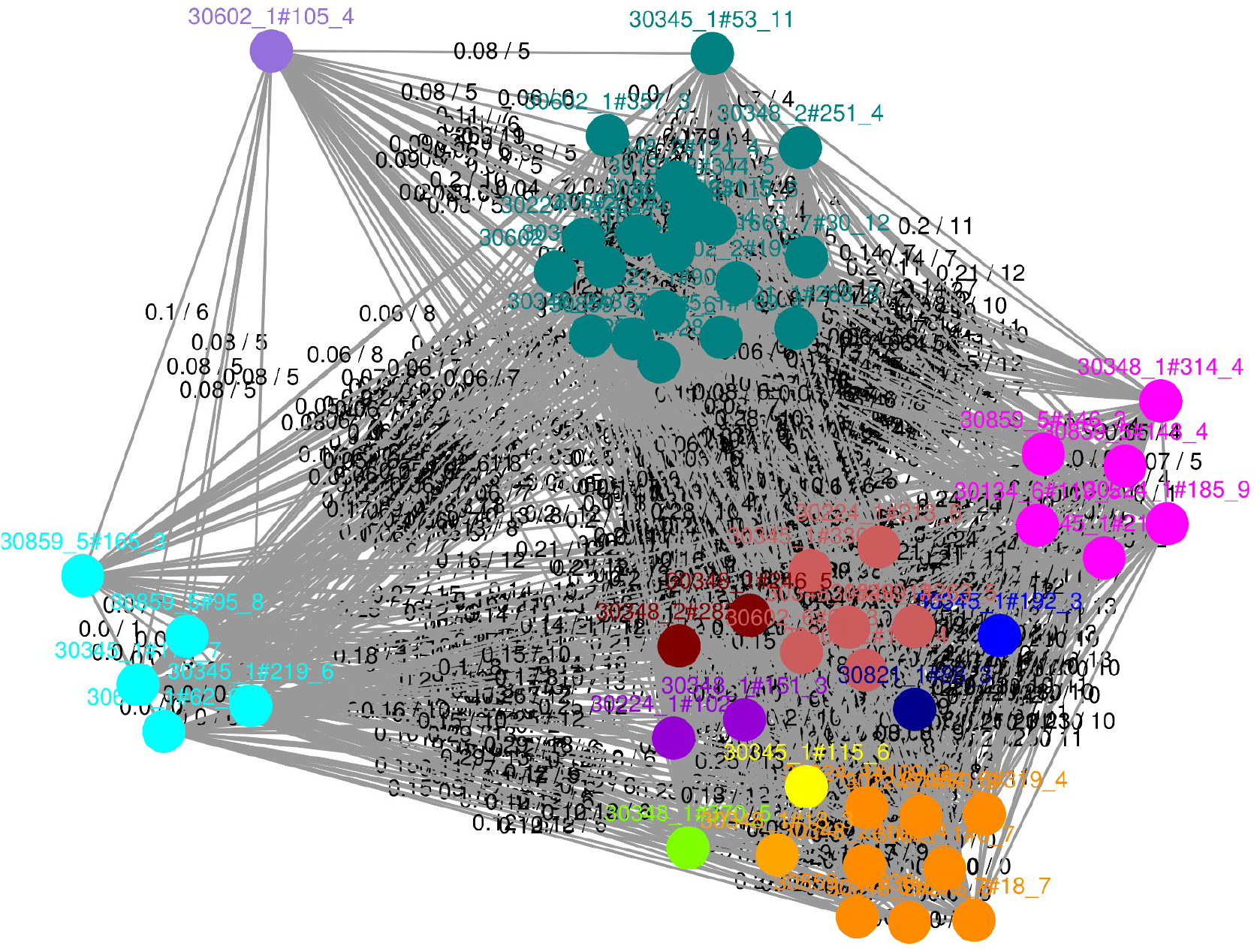
Example of a community where visual inspection shows changing DCJ-Indel threshold would be reasonable. Visualisation of a containment community, where each subcommunity is indicated by the same colour of node, and edges are labelled by containment and DCJ-Indel distances. The turquoise, lilac and green subcommunities have lower containment and DCJ-Indel distances between each other than to the remaining subcommunities. They could be joined by choosing a more permissive DCJ-Indel threshold, if this is desired by the user. This is a community from clustering plasmids from (46).

The containment threshold is by default 0.5, and it is not advisable to make it less strict (i.e. larger), for the following reason: DCJ-Indel is only meaningful if applied to a pair of plasmids that share a reasonable amount of sequence – otherwise, in the extreme, one can change one plasmid into a totally different second plasmid in just 2 DCJ-Indel steps by deleting the first plasmid, and inserting the second plasmid. However, if you are working on short evolutionary scales, it can be sensible to reduce the containment threshold.

In Figure 5, plasmid B connects A to C and D, by being just under the default thresholds. For the pairs A and B, and C and D, there is clearly a recent evolutionary history connecting them, as is apparent from the small containment and DCJ-Indel distances, as well as the alignments in Figure 5. The connection of these two pairs to each other is less obvious, but becomes clearer when looking at the replicons – the largest plasmid, A, has three replicons, while the remaining three only have one, which is also shared with the largest. The other larger plasmid in this subcommunity, B, is differentiated from A by a deletion, where presumably the other two replicons were lost. However, a large part of this deletion actually maps to the two smaller plasmids, C and D. All this indicates that the pair A and B are the result of a fusion event, and are linked by a common ancestor to the other pair. Hence this clustering does reflect a shared evolutionary history, but likely on a longer time scale. So for e.g. studying hospital transmission of plasmids over the course of two years, this clustering is too broad, and it would be sensible to make the containment threshold stricter to filter out these older evolutionary connections. Typically lowering the threshold to 0.2 or 0.3 is enough, and in this case would certainly separate the two more recently related pairs.

**Figure 5:**
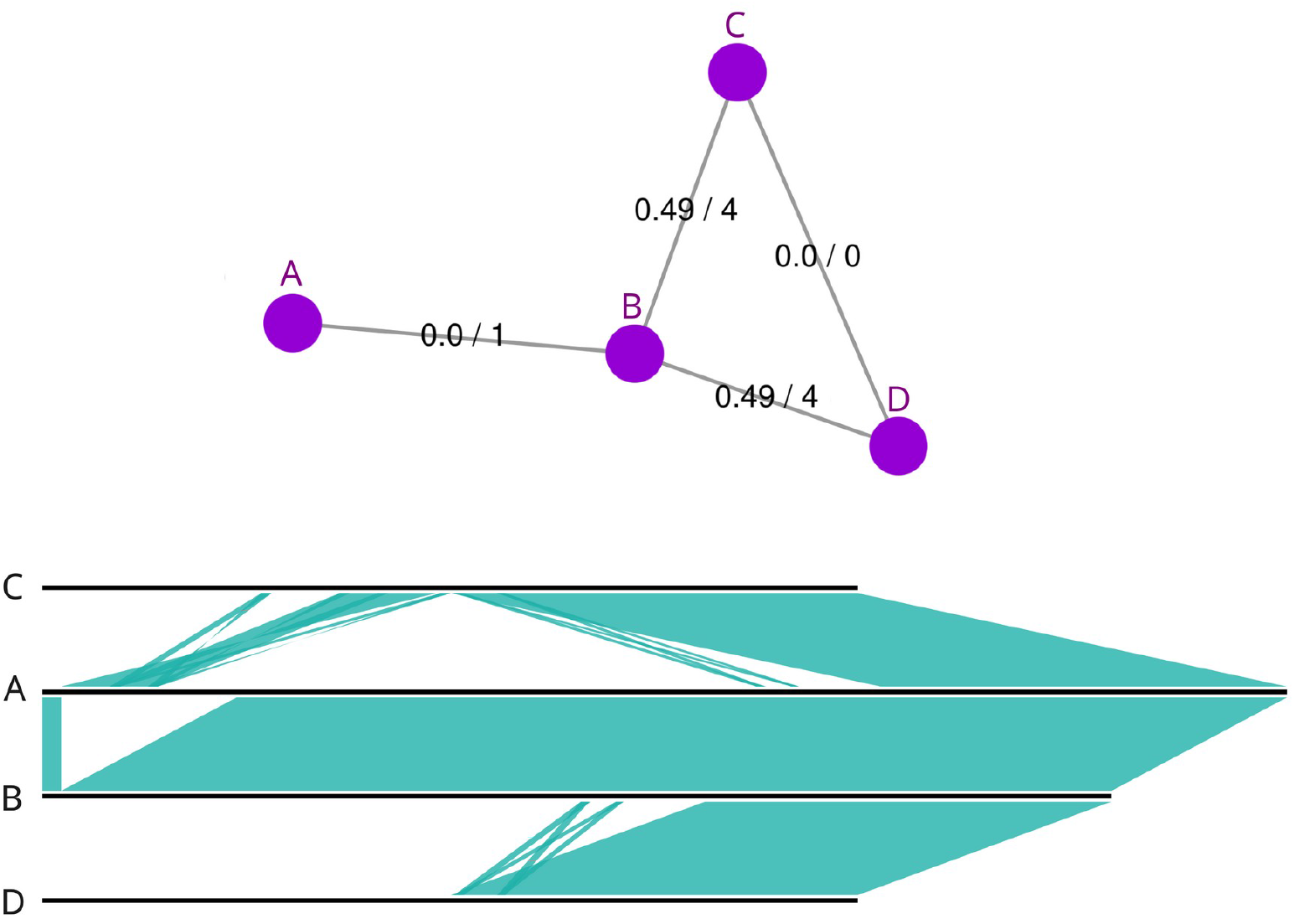
Example for changing containment threshold. The first image is a visualisation of the DCJ-Indel network for this subcommunity of plasmids, while the second is their alignment. The alignment appears to demonstrate a distant but shared common ancestor to all four of these plasmids, which is also supported by replicon typing. However, this relation appears to be too distant to be accounted for in a transmission analysis, so lowering the containment threshold is warranted. This example is from the Addenbrookes hospital dataset analysed in (3, 47).

When the containment and DCJ-Indel thresholds generally seem to produce good subcommunities, aside from pulling in a lot of unrelated plasmids through one or two particular plasmids, then it can be worth increasing the edge density threshold for defining hub plasmids. Hub plasmids are determined based on how many neighbours they have, and how densely connected those neighbours are – the idea is that a hub connects things that are otherwise unrelated. The minimum number of neighbours a plasmid needs to have to be considered a hub is set through the *--bh_connectivity* argument, while the maximal connectivity of its neighbours is controlled through *--bh_neighbours_edge_density*. In Figure 6, the subcommunity produced with the default edge density threshold includes a plasmid that looks like it should be a hub, and is adding a lot of unrelated plasmids to an otherwise tight cluster. An additional indicator that this cluster is not quite right is that it lacks a core genome, and contains many diverse Inc types. Increasing the threshold results in the subcommunity to the right, and it is much cleaner and does not include unrelated plasmids; this could be verified by checking that it does now have a core genome, and the plasmids’ Inc types are mostly shared. A consequence of increasing this threshold is that more plasmids are labelled as hubs, and this can result in smaller subcommunities. However, if you are looking for clusters with a significant and consistent backbone, filtering hub-like plasmids more aggressively is typically beneficial.

**Figure 6:**
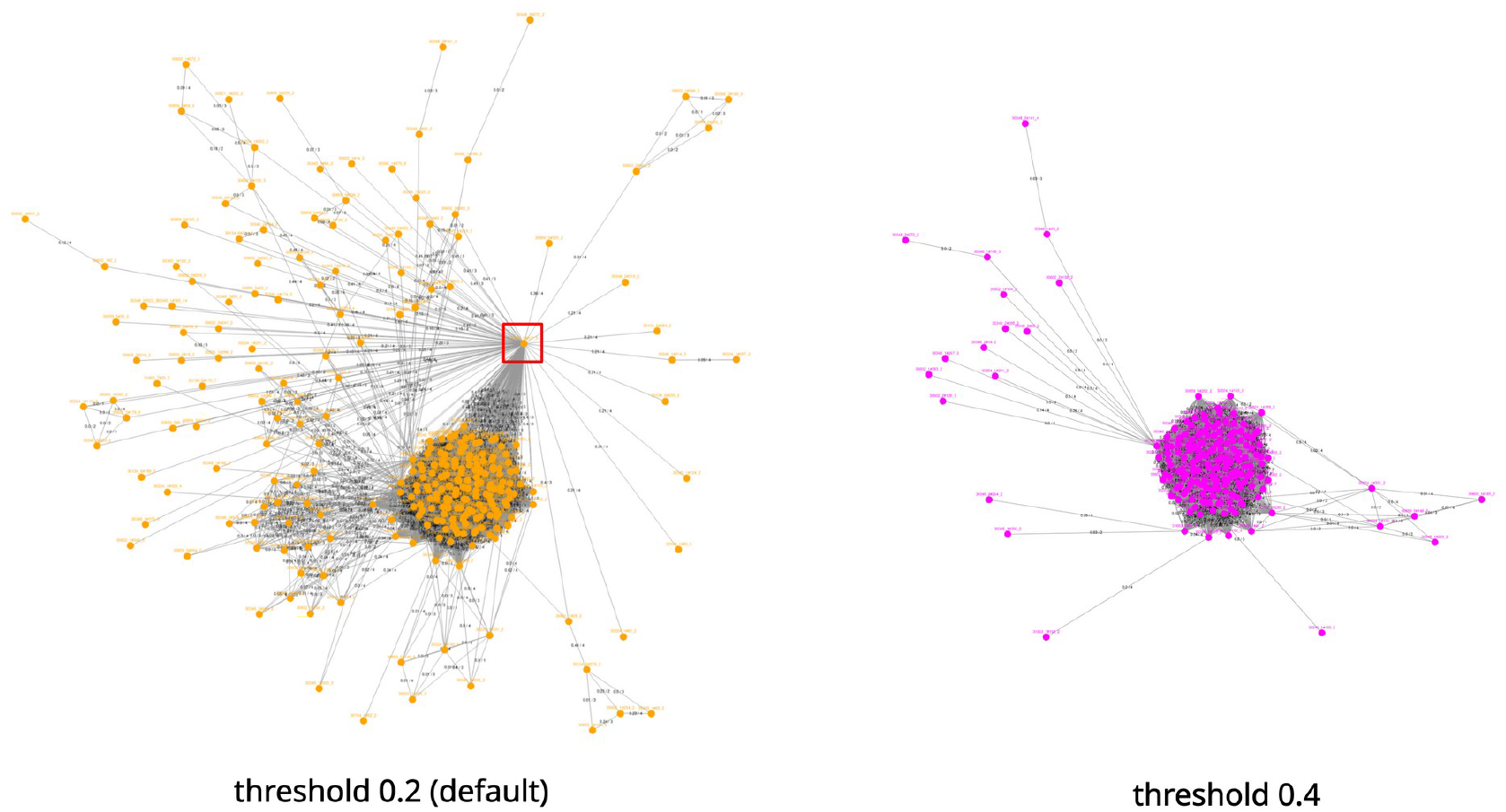
Example for changing edge density threshold for defining a hub. In the subcommunity network on the left, there is a densely connected plasmid whose neighbours are sparsely connected (framed by red square), all of which have been clustered into one subcommunity. The structure of the network, and the lack of common genes between these plasmids, indicates that this was a hub plasmid that was not found when clustering with default thresholds. Increasing the default edge density threshold has resulted in still clustering the same densely connected plasmids as in the previous clustering, but the unrelated plasmids are now removed. These clusters were generated from plasmids from (46).

### 9. Interpreting pling visualisations to identify evolutionary events

#### 9.a) Cointegrates

Broadly speaking, there are two ways to spot cointegrates in the networks: firstly, at community level, by looking at how subcommunities connect to each other; secondly, at subcommunity level, by observing whether or not it looks like there may be further subclusters within a subcommunity. In the first case, in the community visualisation there will be two (or more) subcommunities that connect through a plasmid that is densely connected to both subcommunities, and there are no other connections between the subcommunities. Like in Figure 7, the bridging plasmid will typically be assigned to one of the two subcommunities (generally whichever it has more connections to). This type of topology is an indicator that the bridging plasmid may be a fusion of plasmids from the two subcommunities, and can be confirmed by comparing the lengths, Inc types and/or aligning the bridging plasmid with other plasmids from both subcommunities. For Figure 7 this was confirmed by comparing the alignment, and it is the same example as in Figure 4 in (3).

**Figure 7:**
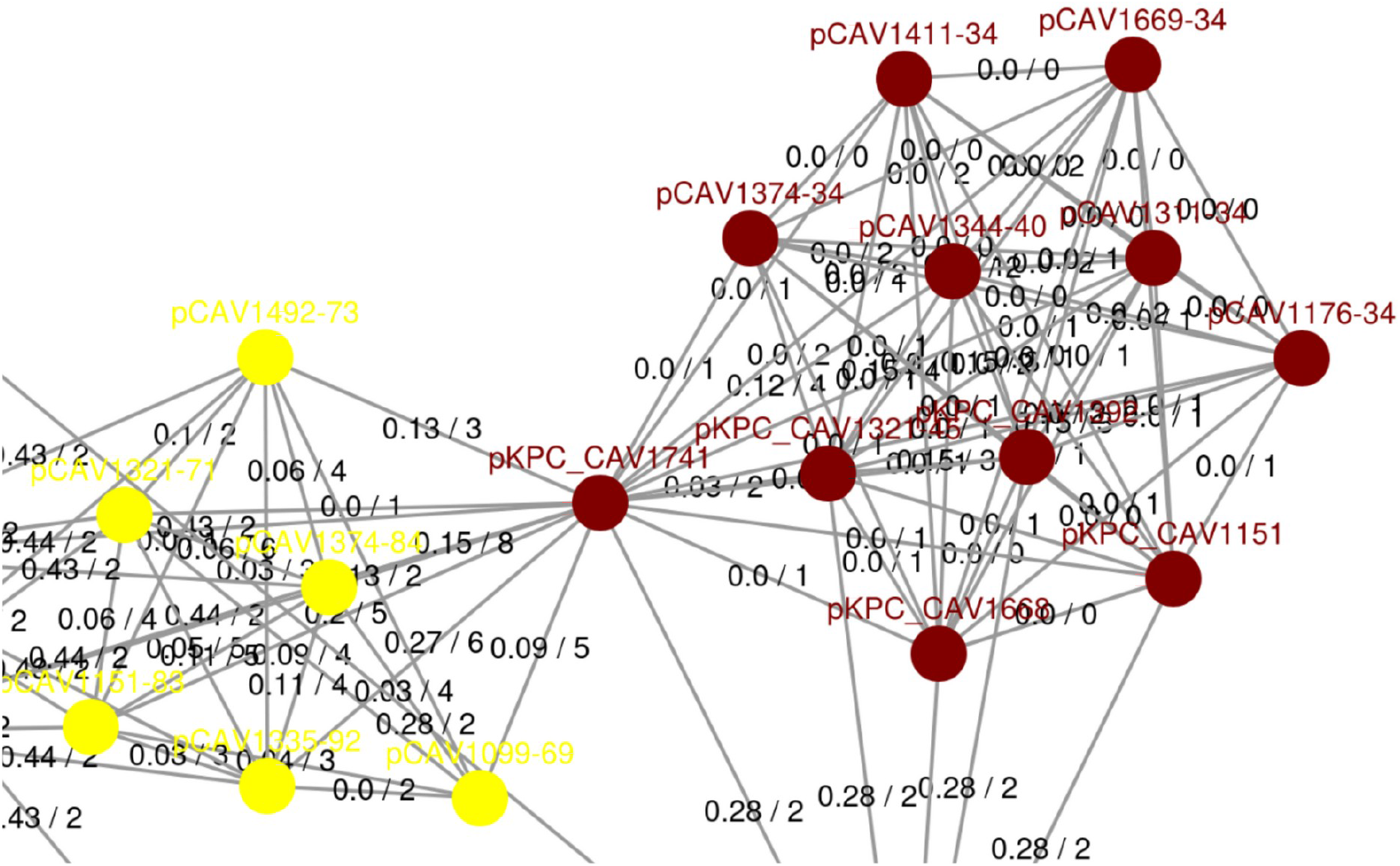
Cointegrate bridging subcommunities. Here is a part of the containment network for the Russian doll plasmid dataset. A single plasmid bridges two subcommunities, and further analysis of alignments between these plasmids revealed it to be a cointegrate, created from the two subcommunities that it bridges. This example was also discussed in (3), and uses data from (21).

There are also situations where parental plasmids and their children will be clustered into a single subcommunity. These kinds of subcommunities typically will have three subgroups, where one bridges the other two, similar to the case discussed above, and as can be seen in Figure 8. Same as before this can be verified through comparing lengths, Inc types, and alignments.

**Figure 8:**
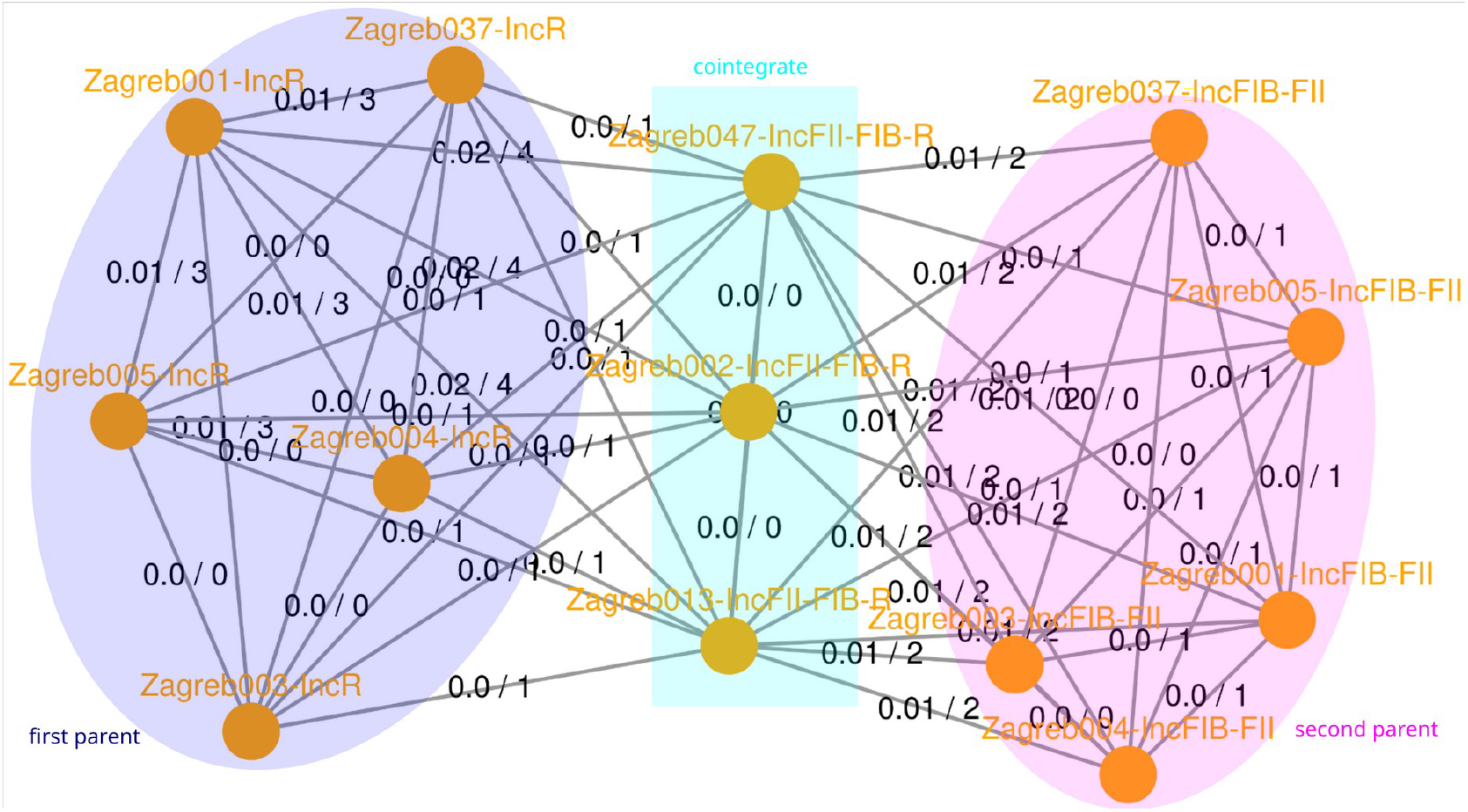
Cointegrates and parental plasmids in a single subcommunity. A DCJ-Indel network for a subcommunity consisting of a cointegrate and its two parental plasmids. The two parents and the cointegrate are each highlighted with a unique colour. The nodes are labelled with both plasmid ID and Inc type – the replicons confirm that this network is showing the outcome of a fusion event. This subcommunity was found in (48).

Another example of network topology caused by cointegration events, is when all plasmids are densely connected in the subcommunity, but there are two subgroups with very small containment distances within them, but distances close to 0.5 between them. This is generally caused by two types of fusion plasmids sharing a single parent; similar to the example discussed in Figure 5.

#### 9.b) Hub plasmids

Hub plasmids are easy to spot as they are denoted by a black star, like for example in Figure 9. They are usually relatively small plasmids that are dominated by a transposable element, which has spread across multiple divergent plasmids. In Figure 9, all the plasmids containing the transposable element have the same containment and DCJ-Indel distances to the hub plasmids, which is often the case with hub plasmids. However, the spread of a transposable element is not the only event that can cause a hub – smaller plasmids that often integrate with other, larger plasmids can become hubs (e.g. this is the most likely origin for the hub in Figure 6), and also hubs can be very large plasmids that are fusions of multiple plasmids, or have accumulated multiple widespread genetic elements. To distinguish between all these cases, examining the lengths and annotating transposons is recommended.

**Figure 9:**
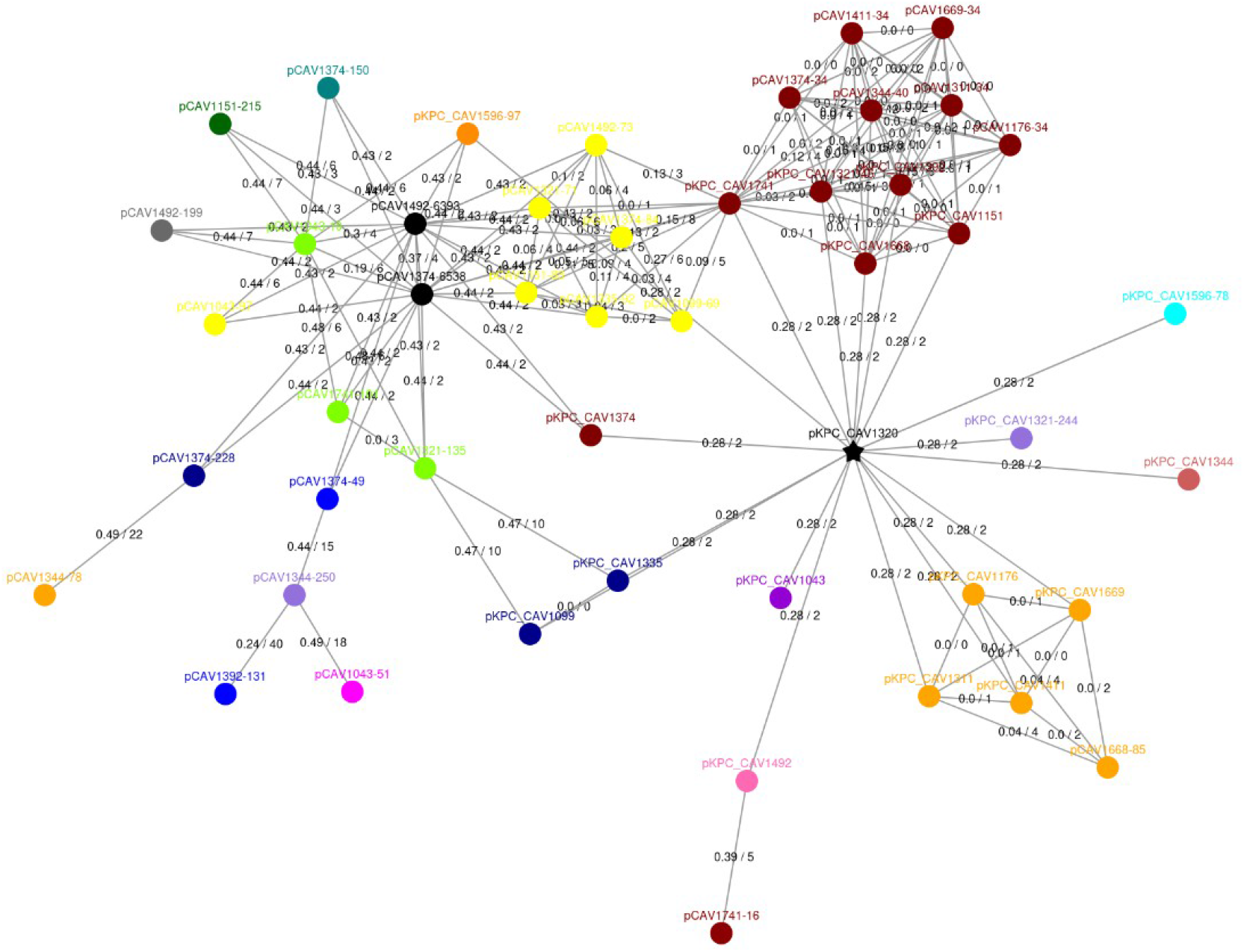
Hub plasmid. This shows the containment network of a plasmid community. The black star denotes a hub plasmid. This is the same hub plasmid as discussed in (3), and is from data in (21).

### 10. Integration with host and SNP information

Once you are satisfied with your clustering, you will likely want to find out how the plasmids are distributed across their bacterial hosts. A simple way to do this is by constructing a tree from the host samples (with e.g. parsnp), and colouring the leaves by which plasmid subcommunities are present in a sample – as in for example in Figure 6 in (3). With this, you can visually judge how plasmids are distributed: if they are being vertically inherited in particular clades, or horizontally spread across different ones.

There are also bioinformatics tools which allow detection of significant clades in the host tree based on presence or absence of a plasmid subcommunity – i.e. tree factorisation approaches like phylofactor or TreeSeg (41, 42). They are both R packages and are available, with documentation, on GitHub. These can be useful on larger datasets, where visually spotting patterns in trees is harder, and also to computationally verify clades identified visually.

A useful analysis is also evaluating the reverse – the distribution of hosts within a plasmid subcommunity. It is possible to construct neighbour joining trees using the DCJ-Indel distances (we suggest quicktree (43)), but this must be done with extreme caution. Firstly, you can always make a hierarchical clustering (tree) of plasmids, but it may not have any phylogenetic meaning, as plasmids do not necessarily have a tree-like evolutionary history. Secondly, DCJ-Indel is defined pairwise and as the minimal number of changes necessary, which means it can skew branch lengths. If several consecutive insertions occur at one position, they will appear to DCJ-Indel as one larger insertion, which will artificially shorten branch lengths.

Finally, there is a pragmatic issue with creating a NJ-tree from the pling distance output, which is that pling only calculates DCJ-Indel distances for plasmid pairs that share sufficient sequence (i.e. are under the containment threshold), so some distances may be missing within a subcommunity. These distances can be missing for awkward technical reasons, which can be circumvented, or because the evolutionary history of these plasmids is fundamentally impossible to (meaningfully) show as a tree. There are three reasons why this can happen:

1. The first case is where a pair of plasmids are separated by a containment distance that is just barely above the containment threshold, and so pling did not calculate their DCJ-Indel distance. If those plasmids are otherwise well connected within the subcommunity, it can be acceptable to calculate a tree.
2. Sometimes sourmash prefiltering falsely excludes a pair due to incorrectly approximating the containment distance; this can happen particularly with small plasmids (<10kb). In this case it is completely safe to calculate DCJ-Indel distances and create a tree.
3. The containment distance is calculated correctly and close to 1 – in this case, the DCJ-Indel distance between this pair will only be 2. This is because for two completely different plasmids, it only takes 2 indel operations to transform one into the other: first by deleting a whole plasmid, and then inserting the other. These pairs will, for example, be found in subcommunities where a fusion has occurred, and both parent plasmids and the child cointegrate are present. The two parents then have nothing in common, and so have containment distance 1. These subcommunities are inherently not tree-like, and cannot be represented by a DCJ-Indel tree.

Generally you will be able to distinguish between these three cases by studying the neighbourhood of a problematic pair on the containment network. If the containment distances are below the threshold between them and all their shared neighbours, then you are likely in the first two cases. If the pair has few or no shared neighbours, and the network looks similar to those discussed in 9a), then you are in the third case.

Currently the simplest way to compute missing DCJ-Indel values is to rerun pling on the subcommunity with the containment threshold set to 1.

When constructing a DCJ-Indel tree is possible, you can colour the leaves by their host, which will allow you to see if the plasmid population structure seems to be influenced by host – e.g., if an outlier on a distant branch has a different host to the rest, that may imply there is host specific adaptation. In Figure 10 is a subcommunity of *E*.*coli* plasmids, each coloured by their host ST, and the tree can clearly be separated into clades on the basis of host ST.

**Figure 10:**
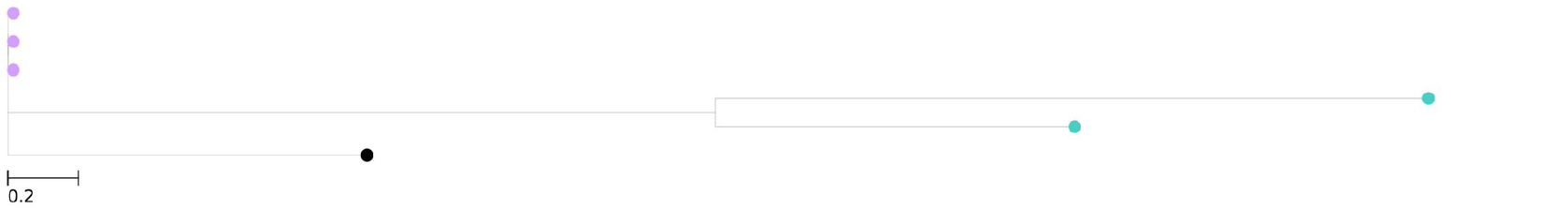
DCJ-Indel neighbour joining tree with host correlated population structure. Shown is a DCJ-Indel neighbour-joining tree of a plasmid subcommunity. Each node is coloured by the ST of the sample the plasmid is from. This visualisation shows that there are greater differences in plasmids from different STs, than within the same. This is a subcommunity found by clustering (46).

As pling subcommunities often have significant and consistent backbones, SNP analysis is feasible. There are two possible ways to build SNP trees – you can use the core genes from ggcaller to build a core gene SNP tree, or alternatively, you can run parsnp, which will find a core genome itself and build a SNP tree from that. Often SNP trees will be very flat, and not very informative, as plasmids accumulate SNPs very slowly, if at all. Nonetheless, it can be useful to generate these trees, since like for DCJ-Indel distances, you can use them to find possible associations between host and plasmid.

## Notes

Here we describe how to use pling on plasmids. However, the underlying method does not explicitly use any plasmid specific information for its clustering, and so can be applied to other genomes. For bacteria, the bottleneck is computation time of the integerisation step, but this can be skipped by providing an integer sequence for each genome upfront. You can use ggcaller to create such integer sequences in terms of genes. We have also extended the tool to cluster genomic regions. For further details, please refer to the pling documentation (44).

## Notes

### Competing Interest Statement

The authors have declared no competing interest.

